# UV inactivation of bacteria and viruses on surfaces: mechanistic insights and testing method comparisons

**DOI:** 10.64898/2026.06.23.734141

**Authors:** Ben Ma, Saba Seyedi, Karl G. Linden

**Affiliations:** Department of Civil, Environmental, and Architectural Engineering, University of Colorado Boulder, 4001 Discovery Dr., Boulder, CO, 80303, United States; Department of Civil and Environmental Engineering, University of Nevada, Reno, 1664 N Virginia St., Reno, NV, 89557, United States; Jacobs Engineering, 1851 Alexander Bell Drive, Reston, VA, 20191, United States

## Abstract

Germicidal UV devices offer a promising solution to mitigate surface-mediated pathogen transmission, providing effective disinfection without material corrosion. This study evaluated the surface inactivation kinetics of two bacteria and two bacteriophages using a low-pressure (LP) mercury UV lamp (254 nm) and a filtered krypton chloride (KrCl*) excimer lamp (222 nm). Three deposition methods (Spray, Spread, and Pipette) and two extraction methods (Swab and Elute) were compared. The UV dose response on surfaces followed a two-region non-linear model due to shielding from dried deposition constituents, primarily through UV absorption. KrCl* excimer exhibited similar bacterial inactivation but slightly lower viral inactivation than LP UV lamp (maximum inactivation ∼ 1 log lower), but its safety profile makes it compelling in occupied spaces. Compared to aqueous conditions, bacteria were more UV sensitive on surfaces, whereas viruses were more resistant. The deposition methods affected the inactivation results, with the Spray method resulting in higher bacteria inactivation. While the extraction methods had limited effect on inactivation efficacy, the Swab method provided higher inactivation detection limits (∼ 2 log higher) and more consistent extraction efficiency. This study provides mechanistic insights into the effects of deposition conditions, UV wavelengths, and microbial characteristics on UV surface disinfection and contributes to standardization of testing methods.

**TOC Graphic:** 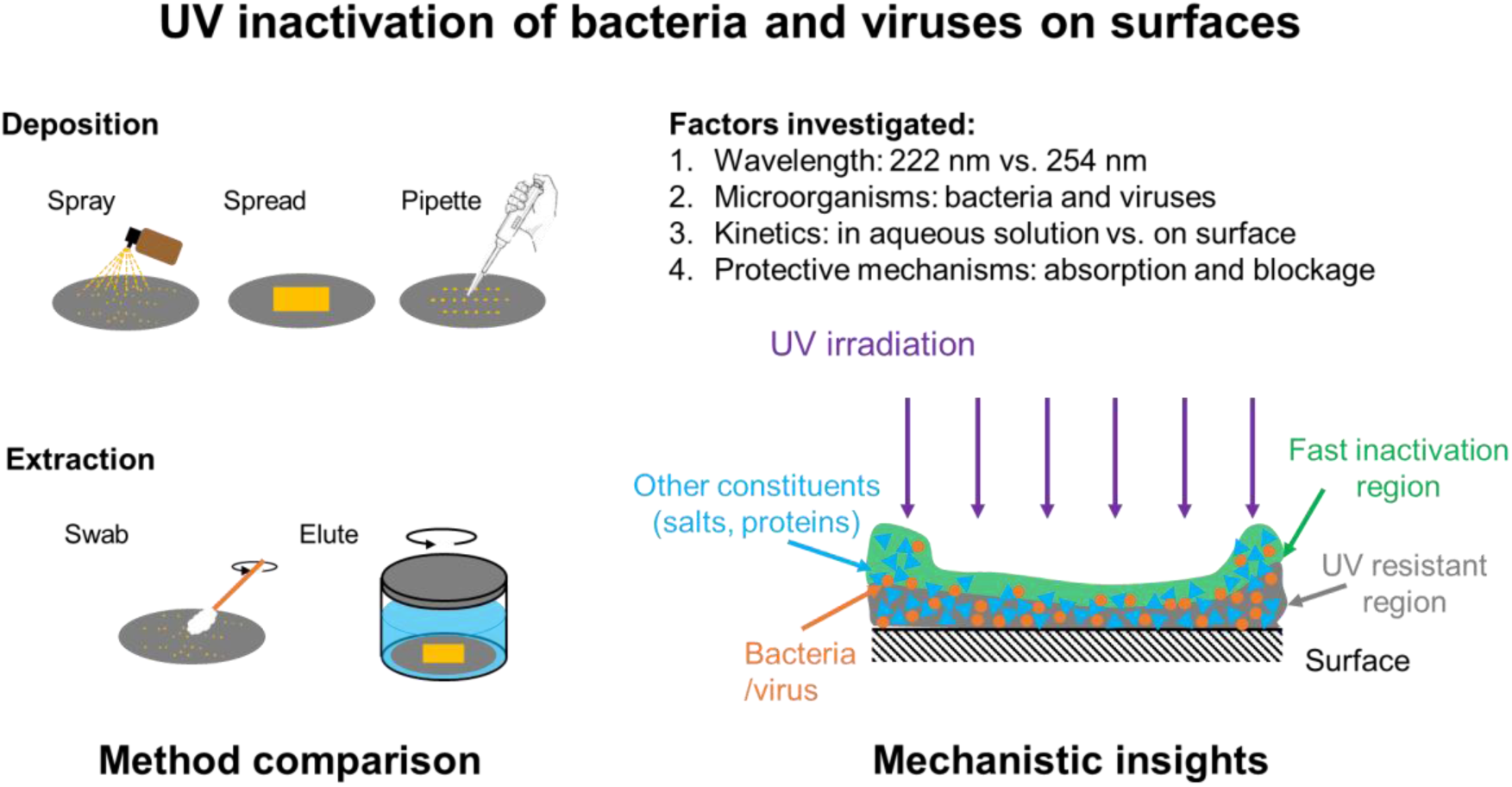

## 1. INTRODUCTION

Pathogen transmission via contaminated surfaces (i.e., fomite transmission) is considered as a major transmission route of various pathogens, including bacteria, viruses, and fungi^1,2^. Considering pathogens may stay viable on surfaces up to several days^3^, this poses serious risks to public health, especially in occupied indoor environments such as healthcare facilities and public transportation systems. Chemical oxidants, such as alcohol, chlorine compounds (e.g., bleach), and hydrogen peroxide, are the most commonly used for surface disinfection^4^. There are, however, several disadvantages^5,6^, including low effectiveness against certain pathogens, intermittent application (can not be applied continuously), material corrosion, and potential adverse health effects such as respiratory irritation and eye and skin damage.

Germicidal ultraviolet (UV) irradiation (200-300 nm) has been proven to be effective for inactivation of all types of pathogens via damage to nucleic acids and proteins^7–9^. Compared to chemical disinfectants, UV irradiation has several advantages for surface disinfection, including high effectiveness and limited-to-no corrosion^10–12^. Conventional UV devices, such as low-pressure (LP) mercury vapor lamps emitting at 254 nm, have been used for surface disinfection in medical and food industries^13^. Their use, however, is typically limited to enclosed configurations (e.g., UV cabinet) and unoccupied spaces due to the risks of 254 nm UV exposure being hazardous to human skin and eyes, including erythema and photokeratitis^14–17^. Emerging UV devices, like the krypton chloride (KrCl*) excimer emitting primarily at 222 nm, can provide effective disinfection performance^18^ and are much safer for human exposure due to strong absorption by the stratum corneum of the skin and the tear film of the eyes, which limits penetration into viable epidermal or corneal cells and reduces tissue damage^19–21^. These features make KrCl* excimer lamps a promising technology providing continuous, in-situ pathogen control on surfaces, especially in high-risk occupied indoor environments.

A few studies investigated the performance of UV devices for pathogen inactivation on surfaces^22–34^. Most studies focused on conventional LP UV device and reported effective inactivation achieved with reasonable UV dose (up to 1-5 log reduction with less than 50 mJ/cm^2^). However, significant variations in the UV inactivation results were observed across studies. Several studies showed that the pathogen deposition method may result in variable UV inactivation^22,29,30,32^, likely due to the shielding effect provided by the deposition mixture. A better understanding of the shielding mechanisms is critical for improving standard test protocols and ensuring consistent and accurate evaluation of UV surface disinfection performance.

In this study, the UV inactivation kinetics of two bacteria and two bacteriophages on surfaces was determined using LP UV and KrCl* excimer lamps. The effects of deposition solution absorbance, deposition methods (Spray, Spread, Pipette), and surface extraction methods (Swab, Elute) on UV surface disinfection performance were evaluated. The UV dose response of bacteria and bacteriophages on surfaces and in aqueous solutions were also compared, providing valuable insights on the mechanisms of UV inactivation and the shielding effects on surfaces.

## 2. MATERIALS AND METHODS

### 2.1. Phage and bacteria solution preparation and enumeration

Two bacteriophages (MS2 coliphage and Phi6) and two bacteria (*Escherichia coli* K-12 and *Staphylococcus aureus*) were used for this investigation. MS2 coliphage (ATCC 15597-B1) and its bacterial host *E. coli* F_amp_ (ATCC 700891) were purchased from the American Type Culture Collection (ATCC). Bacteriophage Phi 6 and its bacterial host *Pseudomonas syringae* were kindly provided by Dr. Michael Fisher’s group at University of North Carolina at Chapel Hill. *Escherichia coli* K-12 (ATCC(R) 29425) and *Staphylococcus aureus* (ATCC 6538) were purchased from ATCC. The detailed methods for virus propagation, purification, and enumeration are included in the supplementary information (SI).

### 2.2. Preparation of virus/bacteria contaminated surface coupons

All experiments were conducted in a biosafety cabinet at room temperature (23 ± 2 °C) with relative humidity (RH) typically ranging from 20% to 35% (uncontrolled). Customized coupons (5 cm in diameter) made of polycarbonate were used for this investigation. Contaminated surfaces were prepared by inoculating virus/bacteria solutions using three different methods described in Table 1 and Fig. 1-A. The Spray-Swab method was developed in this study. The Spread-Elute and Pipette-Elute methods were adopted from American Society for Testing and Materials (ASTM) standard test methods E1153-03^35^ and E3135-18^36^, respectively.

**Figure 1.**
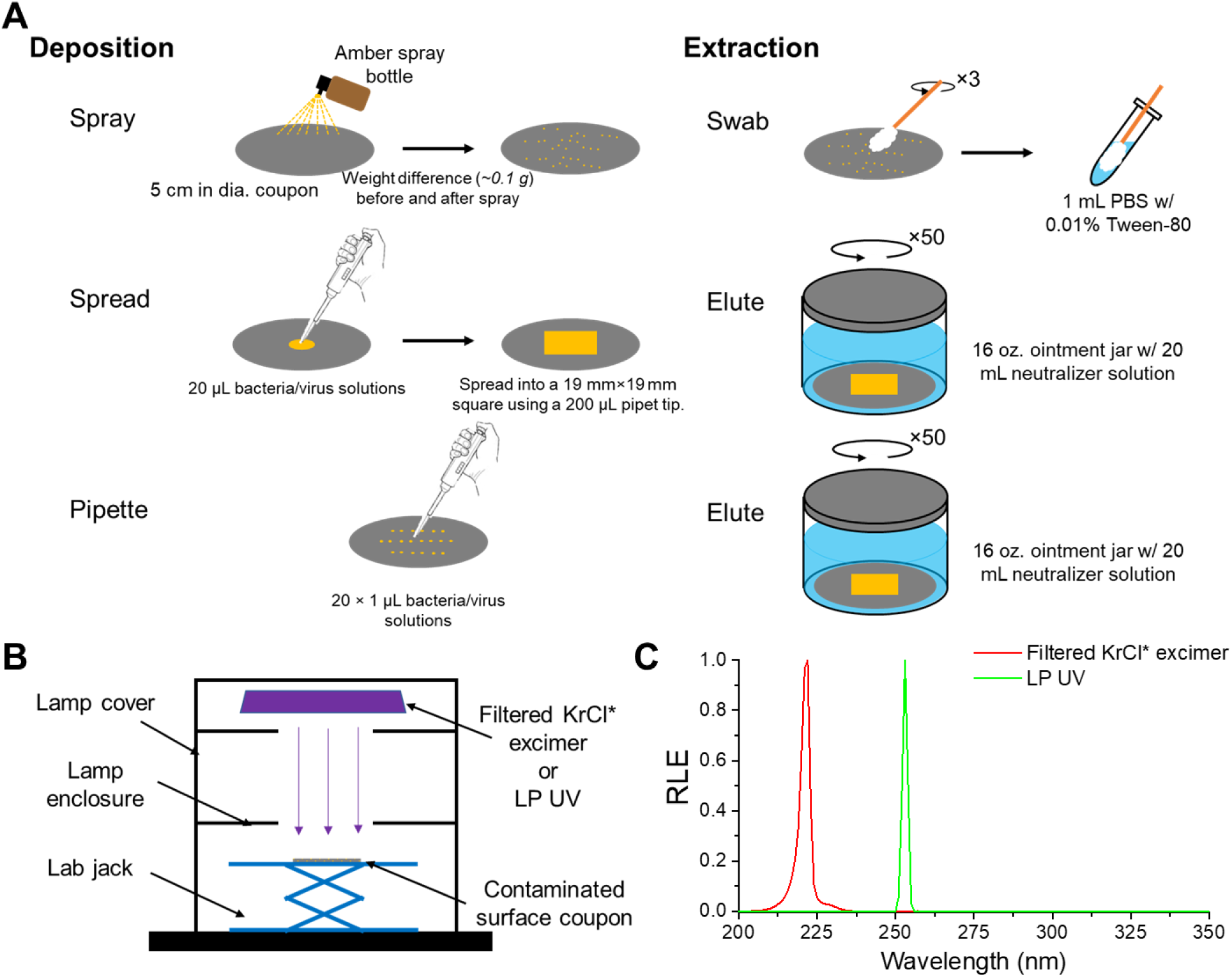
Methods for surface deposition and extraction of tested viruses and bacteria (A), schematic diagram of bench-scale collimated beam apparatus (B), and relative lamp emission (RLE) for filtered KrCl* excimer and LP UV lamp used in this investigation.

**Table 1.**
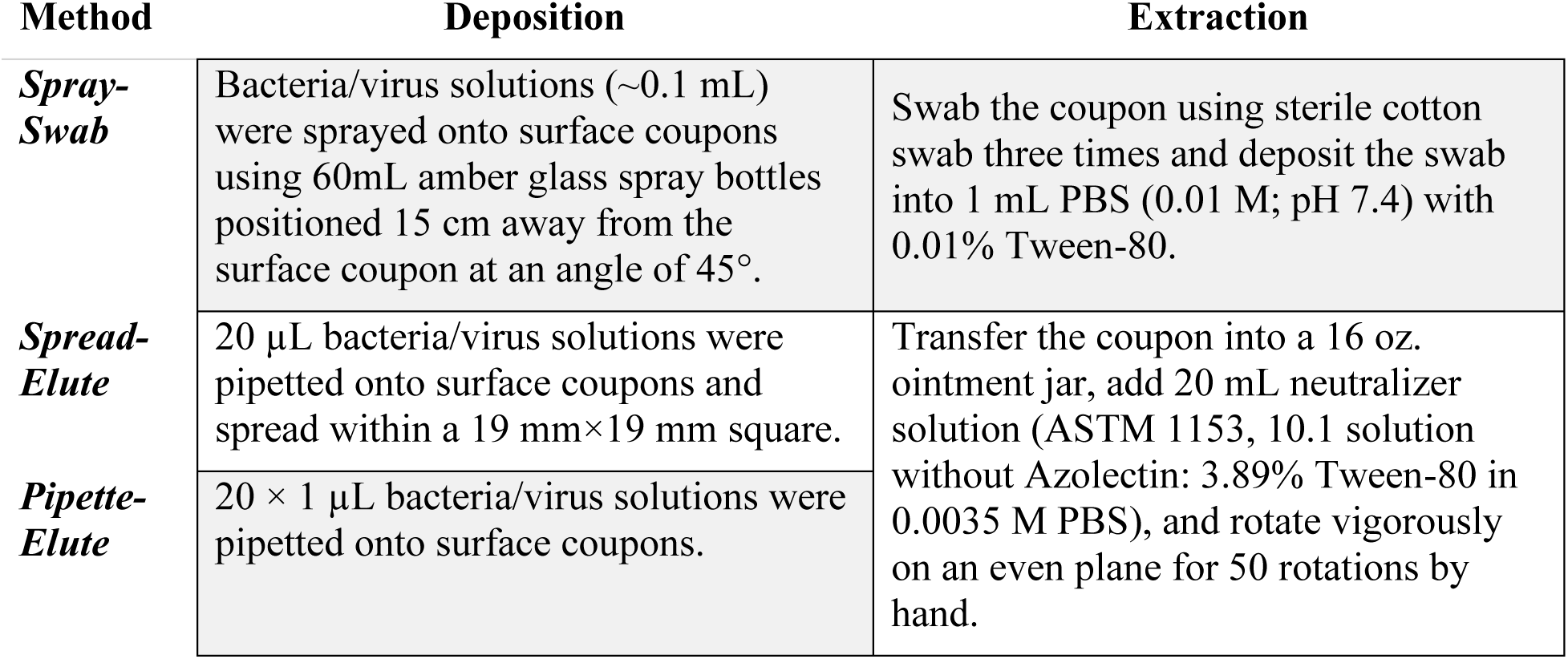
Methods for surface inoculation and extraction of tested viruses and bacteria.

Different deposition solutions were used in this study. Solutions of MS2 and *E. coli* with different absorbance were prepared to study the effects of deposition solution absorbance on UV surface inactivation performance. MS2 stock solutions with no dilution, 10-fold dilution, and 100-fold dilution (in sterile DI) were used for the surface inoculation. The deposition solutions of *E. coli* cells were prepared by resuspending the same amount of washed *E. coli* cell pellets in sterile PBS and cell culture supernatant (collected by centrifugation of cell culture at 3000 g for 5 mins) with no dilution, 2-fold dilution, and 10-fold dilution in DI. The solution absorbance was documented as shown in Fig. 2-C and D. Different solutions of MS2, Phi6, *E. coli*, and *S. aureus* were used to evaluate the difference in all three testing methods and compare the surface disinfection performance between KrCl* and LP UV lamps. The solution preparation details are listed in Table 2. Different dilutions were used to achieve an absorbance of 9 cm^-1^ to 12 cm^-1^ for both 222 nm and 254 nm (i.e., peak emission wavelengths for KrCl* and LP UV lamp, Fig. 1-C), which is similar to the actual human saliva absorbance documented by Barancheshme et al.^32^. Other deposition solution characteristics, such as presence of salts and protein, which may also affect the UV inactivation kinetics, were not considered in this study. The concentrations of the bacteria/virus solutions were measured using the methods described in the SI. The densities of the solutions were also determined by weighing 1 mL of the solution in a microcentrifuge tube (1.0 g/mL for all solutions used).

**Figure 2.**
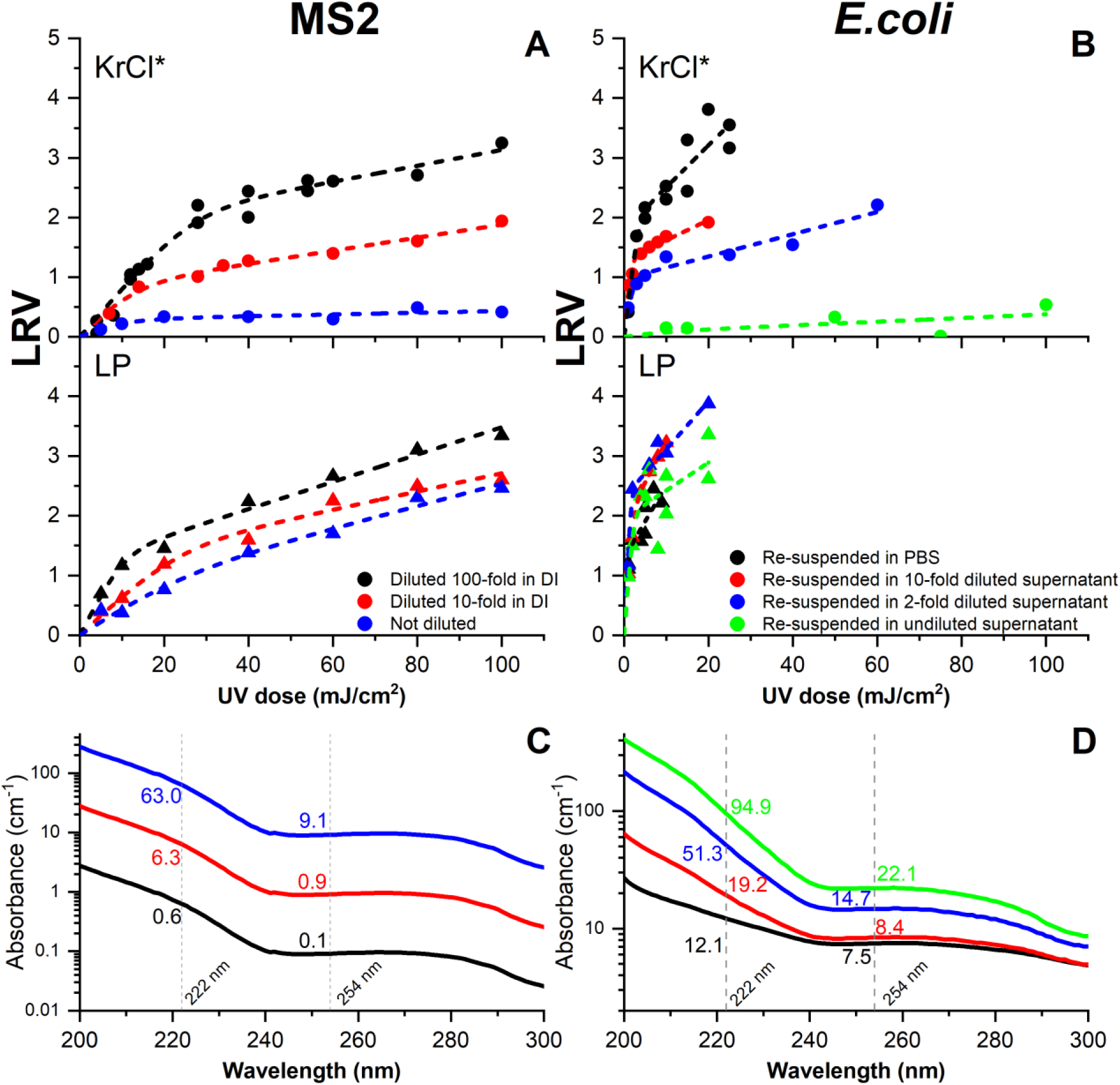
UV inactivation of MS2 (A) and *E. coli* K-12 (B) using the filtered KrCl* excimer (circles) and LP UV (triangles) using the Spray-Swab method. The surface coupons were inoculated with deposition solutions with different absorbance (C and D). Dashed lines represent the non-linear regression models computed based on Eq. 7. UV absorbance of MS2 (C) and *E. coli* K-12 (D) solutions used for surface coupon inoculation, with the UV absorbance (in cm^-1^) at 222 nm and 254 nm labeled. All UV inactivation results in this figure were obtained using the Spray-Swab method.

**Table 2.**
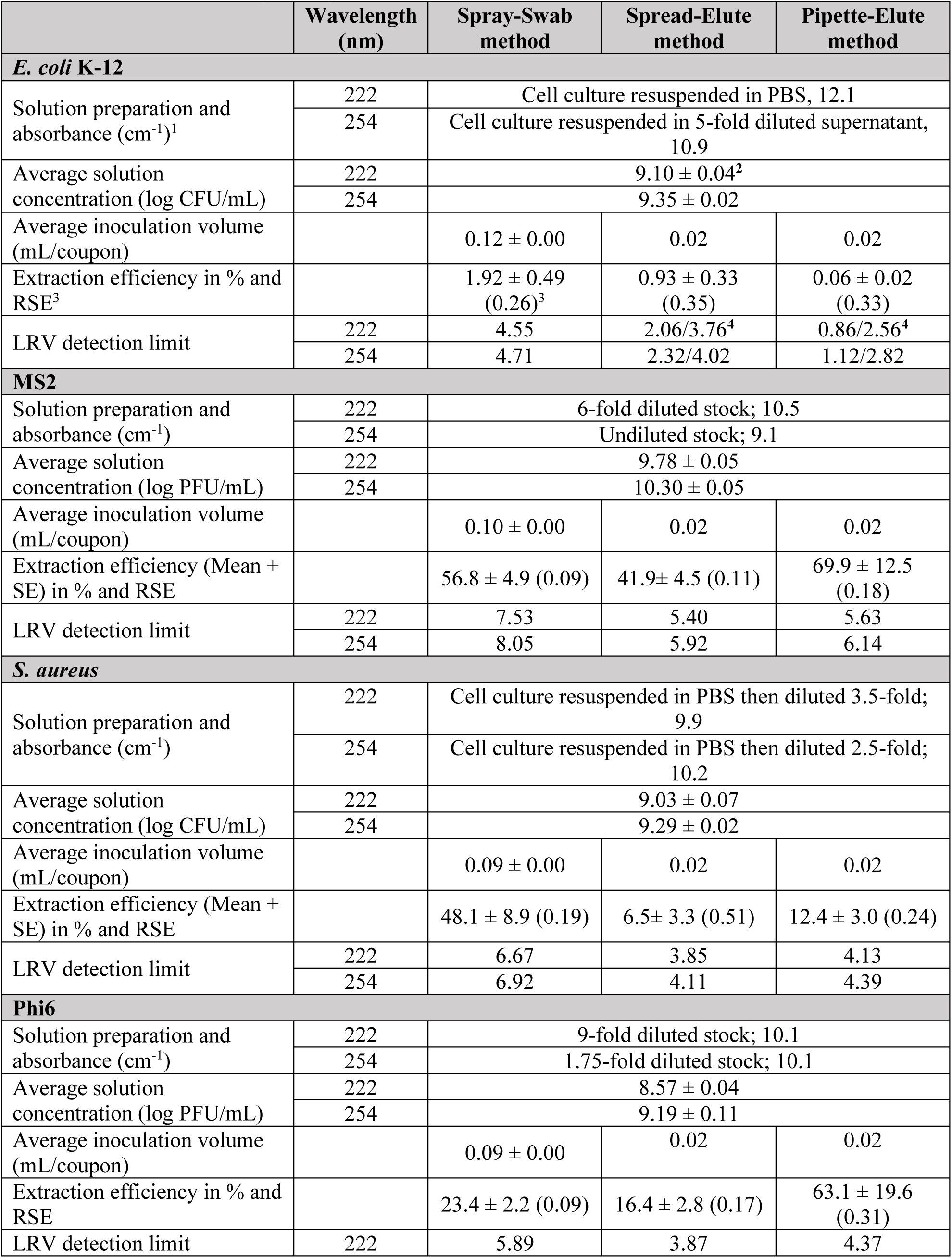
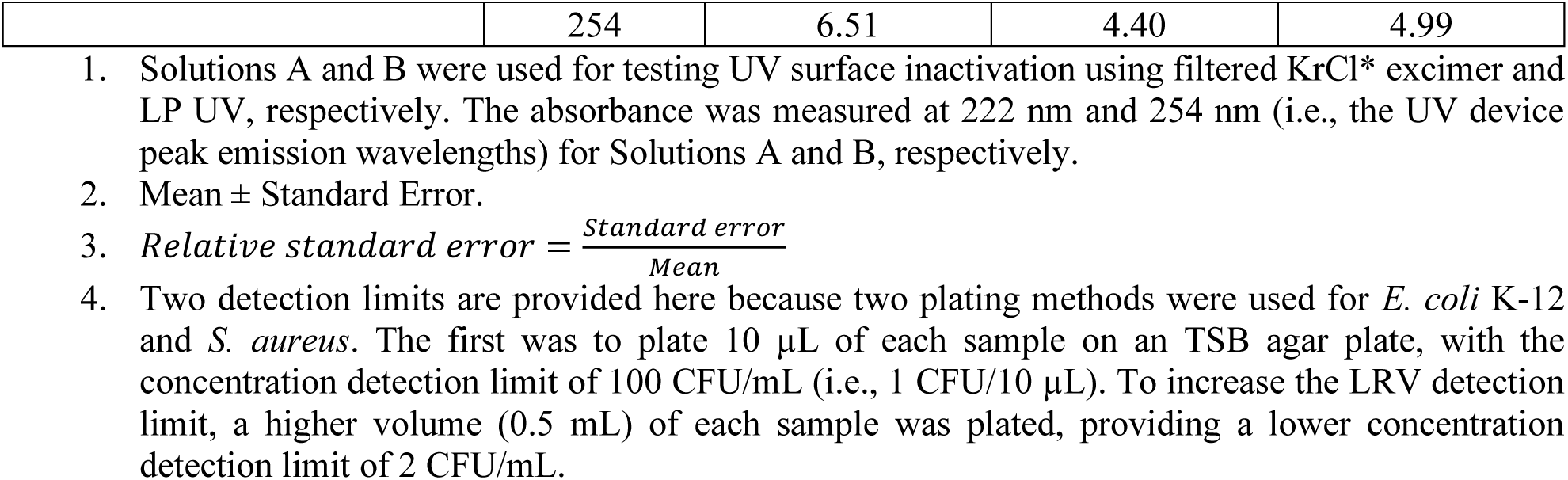
Summary of deposition and extraction information of all three methods.

|  | Wavelength (nm) | Spray-Swab method | Spread-Elute method | Pipette-Elute method |
| --- | --- | --- | --- | --- |
| ***E. coli* K-12** |  |  |  |  |
| Solution preparation and absorbance (cm<sup>-1</sup>)<sup>1</sup> | 222 | Cell culture resuspended in PBS, 12.1 |  |  |
|  | 254 | Cell culture resuspended in 5-fold diluted supernatant, 10.9 |  |  |
| Average solution concentration (log CFU/mL) | 222 | 9.10 ± 0.04<sup>2</sup> |  |  |
|  | 254 | 9.35 ± 0.02 |  |  |
| Average inoculation volume (mL/coupon) |  | 0.12 ± 0.00 | 0.02 | 0.02 |
| Extraction efficiency in % and RSE<sup>3</sup> |  | 1.92 ± 0.49 (0.26)<sup>3</sup> | 0.93 ± 0.33 (0.35) | 0.06 ± 0.02 (0.33) |
| LRV detection limit | 222 | 4.55 | 2.06/3.76<sup>4</sup> | 0.86/2.56<sup>4</sup> |
|  | 254 | 4.71 | 2.32/4.02 | 1.12/2.82 |
| **MS2** |  |  |  |  |
| Solution preparation and absorbance (cm<sup>-1</sup>) | 222 | 6-fold diluted stock; 10.5 |  |  |
|  | 254 | Undiluted stock; 9.1 |  |  |
| Average solution concentration (log PFU/mL) | 222 | 9.78 ± 0.05 |  |  |
|  | 254 | 10.30 ± 0.05 |  |  |
| Average inoculation volume (mL/coupon) |  | 0.10 ± 0.00 | 0.02 | 0.02 |
| Extraction efficiency (Mean + SE) in % and RSE |  | 56.8 ± 4.9 (0.09) | 41.9 ± 4.5 (0.11) | 69.9 ± 12.5 (0.18) |
| LRV detection limit | 222 | 7.53 | 5.40 | 5.63 |
|  | 254 | 8.05 | 5.92 | 6.14 |
| ***S. aureus*** |  |  |  |  |
| Solution preparation and absorbance (cm<sup>-1</sup>) | 222 | Cell culture resuspended in PBS then diluted 3.5-fold; 9.9 |  |  |
|  | 254 | Cell culture resuspended in PBS then diluted 2.5-fold; 10.2 |  |  |
| Average solution concentration (log CFU/mL) | 222 | 9.03 ± 0.07 |  |  |
|  | 254 | 9.29 ± 0.02 |  |  |
| Average inoculation volume (mL/coupon) |  | 0.09 ± 0.00 | 0.02 | 0.02 |
| Extraction efficiency (Mean + SE) in % and RSE |  | 48.1 ± 8.9 (0.19) | 6.5 ± 3.3 (0.51) | 12.4 ± 3.0 (0.24) |
| LRV detection limit | 222 | 6.67 | 3.85 | 4.13 |
|  | 254 | 6.92 | 4.11 | 4.39 |
| **Phi6** |  |  |  |  |
| Solution preparation and absorbance (cm<sup>-1</sup>) | 222 | 9-fold diluted stock; 10.1 |  |  |
|  | 254 | 1.75-fold diluted stock; 10.1 |  |  |
| Average solution concentration (log PFU/mL) | 222 | 8.57 ± 0.04 |  |  |
|  | 254 | 9.19 ± 0.11 |  |  |
| Average inoculation volume (mL/coupon) |  | 0.09 ± 0.00 | 0.02 | 0.02 |
| Extraction efficiency in % and RSE |  | 23.4 ± 2.2 (0.09) | 16.4 ± 2.8 (0.17) | 63.1 ± 19.6 (0.31) |
| LRV detection limit | 222 | 5.89 | 3.87 | 4.37 |
|  | 254 | 6.51 | 4.40 | 4.99 |

1. Solutions A and B were used for testing UV surface inactivation using filtered KrCl\* excimer and LP UV, respectively. The absorbance was measured at 222 nm and 254 nm (i.e., the UV device peak emission wavelengths) for Solutions A and B, respectively. 2. Mean ± Standard Error. 3. $Relative\ standard\ error = \frac{Standard\ error}{Mean}$ 4. Two detection limits are provided here because two plating methods were used for *E. coli* K-12 and *S. aureus*. The first was to plate 10 μL of each sample on an TSB agar plate, with the concentration detection limit of 100 CFU/mL (i.e., 1 CFU/10 μL). To increase the LRV detection limit, a higher volume (0.5 mL) of each sample was plated, providing a lower concentration detection limit of 2 CFU/mL.

For Spray-Swab method, the coupons were weighed before and right after (within 10 seconds) the inoculation and the virus/bacteria load on each coupon was determined using Eq. 1:

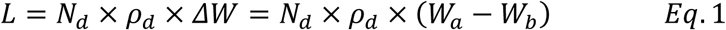

where *L* is the virus/bacteria load in PFU (MS2 or Phi6) or CFU (*E. coli* or *S. aureus*) per coupon, *N_d_* is the concentration of the virus/bacteria solution, *ρ_d_* is the density of the solution, *ΔW* is the weight difference after and before a deposition, *W_a_* and *W_b_* are the weights of the coupons right after and before a deposition.

The virus/bacteria load on each coupon using Spread-Elute and Pipette-Elute methods were calculated as:

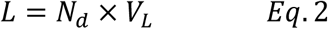

where *V_L_* is the volume of virus/bacteria solutions inoculated on the surface coupon (i.e., 20 µL for Methods 2 and 3).

### 2.3. UV exposure on surfaces

The UV lamps were set up in a bench-scale collimated beam apparatus (Fig. 1-B) as described by Bolton and Linden^37^. Normalized emission spectra for these UV lamps as used in the experiments (Fig. 1-B) were measured using a calibrated Maya 2000 Pro spectrometer (Ocean Insight, Dunedin, FL). Two UV sources were used in this investigation: a KrCl* excimer lamp with a bandpass filter emitting at 222 nm (USHIO, Cypress, CA, USA) and a conventional LP mercury lamp emitting at 254 nm. UV exposure experiments on surfaces were performed according to a standard protocol by Bolton and Linden^37^. After deposition, surface coupons were allowed to dry under sterile laminar flow in a biosafety cabinet for at least 1 hour at room temperature and then placed at the center of the beam on a lab jack in the collimated beam apparatus. The UV incident irradiance at the sample surface was measured using a calibrated radiometer (ILT-2400, International Light Technologies, Inc., Peabody, MA, USA). UV exposure time for each sample was calculated using the target UV fluences for unweighted emissions between 200 nm to 400 nm according to a protocol by Linden and Darby^38^ and Bolton and Linden^37^. Average UV fluence across the coupon surface was calculated by including corrections for radiometer detector sensitivity correction across lamp emission spectra (i.e., lamp correction factor) and non-uniformity of incident irradiance across the sample surface (i.e., petri factor). The water factor, accounting for solution absorbance, was omitted. Coupons were exposed under UV one by one in the collimated beam apparatus, with the order of exposure time randomized to minimize any potential time-dependent bias.

### 2.4. Sample extraction

Bacteria and viruses from contaminated coupons without and after UV exposure were extracted using methods described in Table 1 and Fig. 1-A. The concentrations of bacteria and viruses in the extraction solution were measured using the method described above. The extraction efficiencies of test microorganisms were determined by comparing the amount of microorganisms inoculated onto the coupons and the amount of microorganisms extracted from the coupons without UV exposure, considering both ambient microbial decay (i.e., decay of bacteria/viruses on surfaces over the course of drying and UV exposure experiments) and losses in the extraction process:

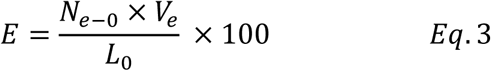

where *E* is the extraction efficiency in %, *N_e-0_* is the concentration of microorganisms in the extraction samples without UV exposure in PFU/mL (MS2 or Phi6) or CFU/mL (*E. coli* or *S. aureus*), *V_e_* is the volume of extraction solution (1 mL for Spray-Swab method and 20 mL for Spread-Elute and Pipette-Elute methods), *L_0_* is the number of test microorganisms inoculated on the surface coupons without UV exposure, calculated using Eq. 1 (Spray-Swab method) or Eq. 2 (Spread-Elute and Pipette-Elute methods). Three control samples (no UV exposure) were collected throughout each UV exposure experiment after the 1-hour drying period, with one control sample collected at the beginning, one in the middle of all UV exposure tests, one right after all UV exposure tests were done. No significant trend in the control sample concentrations (i.e., increase or decay over time) was observed. An average *E* value was calculated from triplicate control samples for each UV exposure experiment.

The UV inactivation performance was calculated as the log reduction value (LRV) taken from the average concentrations of the test microorganisms in the extraction samples for the control (no UV treatment) and after a UV exposure. For Spray-Swab method, the concentrations were normalized by the number of microorganisms inoculated:

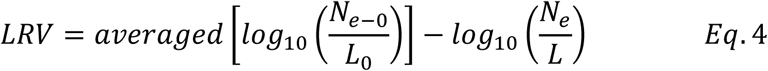

where *N_e-0_* is the concentration of microorganisms in the extraction samples after a UV exposure in PFU/mL (MS2 or Phi6) or CFU/mL (*E. coli* or *S. aureus*) and *L* is the number of microorganisms inoculated on the surface coupon.

For Spread-Elute and Pipette-Elute methods, considering the same volume were inoculated on each coupon (i.e., *L_0_* = *L*), the UV inactivation was calculated as:

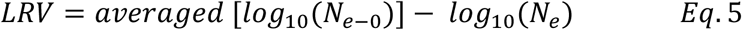

The LRV detection limit for each extraction method was calculated using Equation 6.

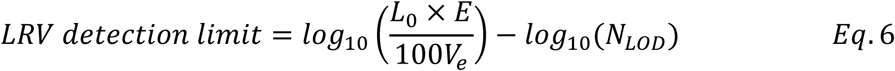

where *N_LOD_* is the limit of detection of test microorganisms, which typically is 1 CFU/10 µL for *E. coli* and *S. aureus* (1 colony in 10 µL samples were plated for enumeration) or 1 PFU/100 µL for MS2 and Phi6 (1 plaque in 100 µL samples were plated for enumeration).

### 2.5. Model for UV dose response on surfaces

The UV dose response of all tested microorganisms on surfaces was evaluated using a two-region non-linear model adopted from Geeraerd and Van Impe inactivation model^39^ and a previous study by Gibson et al.^23^. The dose response curve is divided into two regions: an inactivation region, where the microorganisms were inactivated following a pseudo-first-order kinetics, and a resistant region, where the microorganisms were inactivated at a lower rate than those in the inactivation region. The model equation can be expressed as:

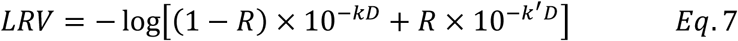

where *D* is UV fluence in mJ/cm^2^ between 200 nm to 400 nm, *k* is the pseudo-first-order inactivation rate constant (in cm^2^/mJ) for the microorganisms in the inactivation region, *k’* is the pseudo-first-order inactivation rate constant (in cm^2^/mJ) for the microorganisms in the resistant region, and *R* is the fraction of the microorganisms in the resistant region. The derivation of Eq. 7 from the original inactivation model is shown in the SI. At least seven data points were included in each UV dose response curve.

The mean and standard error (SE) of *k*, *k’*, and *R* in Eq. 7 and the regression coefficient were determined using the ‘non-linear curve fitting’ function (based on Orthogonal Distance Regression algorithm) in OriginPro 2025 by fitting the observed *LRV* and *D*. The following constraints were applied in the fitting analysis: (1) 0 ≤ *k’* ≤ *k*, (2) 0 ≤ *R* ≤ 1.

### 2.6. Estimation of UV dose response in aqueous solutions

The UV dose responses of all test microorganisms in the aqueous solutions were estimated using kinetics data from previous studies^18^ and compared to the UV dose responses on surfaces. The estimation was conducted based on the aqueous solution having the same UV absorbance and depth as the inoculation solution used for surface disinfection tests. Considering the kinetics data in previous studies were reported using the average UV dose across the exposed sample including the effect of UV absorption, a correction factor was applied to the UV dose on surfaces to include the impact of UV absorbance by the deposition solutions according to the calculation by Bolton and Linden (2003)^37^:

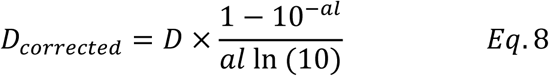

where *D* is UV fluence in mJ/cm^2^ between 200 nm to 400 nm, *D_corrected_* is the UV dose after applying the correction factor, a is the absorbance in cm^-1^, and *l* is the average thickness of inoculation solutions on the surface coupons (average 5.5×10^-3^ cm and 1.6×10^-2^ cm for the Spread-Elute and Pipette-Elute methods, respectively. See SI for calculation details). The UV dose response estimation was only conducted for the Spread-Elute and Pipette-Elute methods but not for the Spray-Swab method due to the difficulty of determining the average depth of deposition solution on surfaces.

### 2.7. Statistical analyses

Two-way ANOVA was used to determine the effect of deposition solution absorbance on UV dose response curve, by setting the UV dose and absorbance as the independent variables. Kendall’s Tau correlation was used to determine if there was a significant correlation between the deposition solution absorbance and the fitting parameters (i.e., *k*, *k’*, and *R* in Eq. 7). Paired t tests were used to determine if there was a significant difference in the UV dose response curves due to using different testing methods (Table 1) or UV devices (KrCl* excimer and LP UV).

## 3. RESULTS AND DISCUSSION

### UV inactivation of bacteria and viruses on surfaces

Both KrCl* excimer and LP UV lamp can effectively inactivate *E. coli* and MS2 on surfaces, providing maximum LRVs of 3.2 and 3.3 at 100 mJ/cm^2^ for MS2, respectively, and 3.6 and 3.9 at 20 mJ/cm^2^ for *E. coli*, respectively (Fig. 2-A and B). Different from the linear UV dose response curve typically observed in aqueous solution (i.e., pseudo-first-order inactivation kinetics with a constant inactivation rate), the UV dose response curve on surfaces exhibited two regions: a fast inactivation region at the lower UV dose range, followed by a resistant region with a lower inactivation rate at higher UV doses. This observation is further supported by the two-region non-linear regression analyses as described in Eq. 7, showing good representation of the UV dose response with regression coefficients (R^2^) greater than 0.9 for most conditions (Table S1). The deviation from linear kinetics on surfaces is likely attributed to physical shielding effects of dried constituents from the deposition solution, such as proteins and salts. The test microorganisms were mixed with these constituents in the dried deposits on surfaces. The microorganisms distributed in the top layer of the dried deposits were likely less protected compared to the bottom layer, resulting in two inactivation regions with different inactivation rates (Fig. 3). Another possible explanation could be due to the “coffee ring effect”, which a greater amount of the deposits was developed along the perimeter of the droplets inoculated on surfaces^40^. The physical shielding effect in the “coffee ring” zone was likely to be greater than other areas, leading to a lower UV inactivation of the microorganisms (Fig. 3). Similar UV dose response on surfaces were also reported in previous studies by Barancheshme et al. and Gibson et al. ^23,32^. Ratliff et al.^22^, however, observed a linear relationship between the LRV and log-scale UV dose, which could also be described using the two-region kinetics model presented in this study.

**Figure 3.**
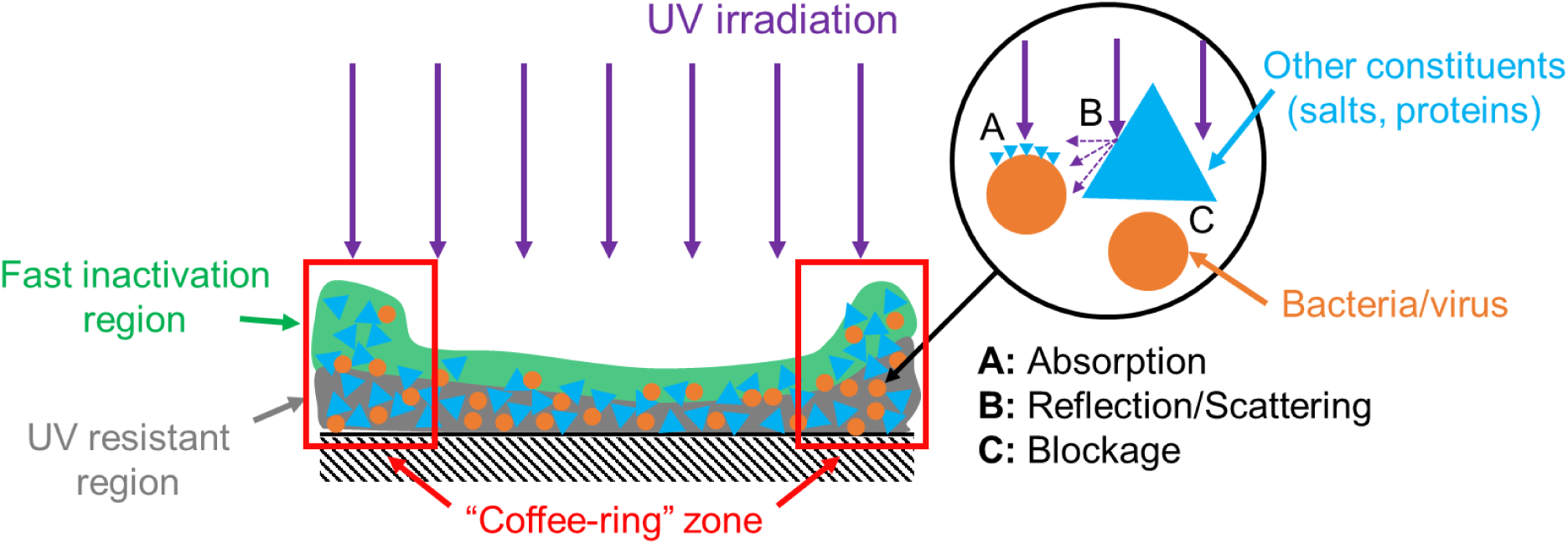
Physical shielding effects of dried deposits on UV surface disinfection.

### Effect of deposition solution absorbance on UV surface inactivation

The UV surface inactivation efficiencies decreased with deposition solution UV absorbance for both KrCl* excimer and LP UV lamp (Fig. 2). For MS2, the observed maximum LRV (at 100 mJ/cm^2^) decreased from 3.2 to 0.4 using the KrCl* excimer and from 3.3 to 2.5 using the LP UV lamp, with the deposition solution absorbance increasing from 0.6 cm^-1^ to 63 cm^-1^ at 222 nm and from 0.1 cm^-1^ to 9.1 cm^-1^ at 254 nm, respectively. For *E. coli*, the maximum LRV decreased from 3.5 to 0.5 using the KrCl* excimer, whereas only a marginal reduction in LRV was observed using the LP UV lamp with UV doses of 0 to 20 mJ/cm^2^, likely due to relatively minimal differences in solution absorbance (less than 3-fold: 7.5 cm^-1^ to 22.1 cm^-1^). This observation is further supported by the regression analyses. Negative correlations between the absorbance and inactivation rate constants (i.e., Kendall correlation coefficient < 0; Figure 4 and Table S2) were observed in both fast inactivation and resistant regions (*k* and *k’*, respectively). Positive correlations between the absorbance and the fraction of microorganisms in the resistant region (*R*) were observed in most experimental conditions except for *E. coli* inactivation using LP UV lamp. This was expected because higher UV absorbance was associated with higher concentrations of proteins and salts in the solution, according to the Beer-Lambert Law (i.e., light absorbance is directly proportional to the solution’s concentration). Upon evaporation, these constituents form more deposits on surfaces, providing greater physical shielding effects (lower *k* and *k’* values) and higher portion of microorganisms that were shielded by the deposits (higher *R* values). It should be noted that some fitting parameters (*k*, *k’*, and *R*) exhibited relatively large standard errors, and limited statistical significance was observed in the correction analysis (Table S2) due to the small sample size (less than 5 absorbance tested in each condition). Previous studies^22,32^ also indicated that deposition solution absorbance may affect the UV surface disinfection performance, highlighting the need to consider deposition solution absorbance in UV surface disinfection tests.

**Figure 4.**
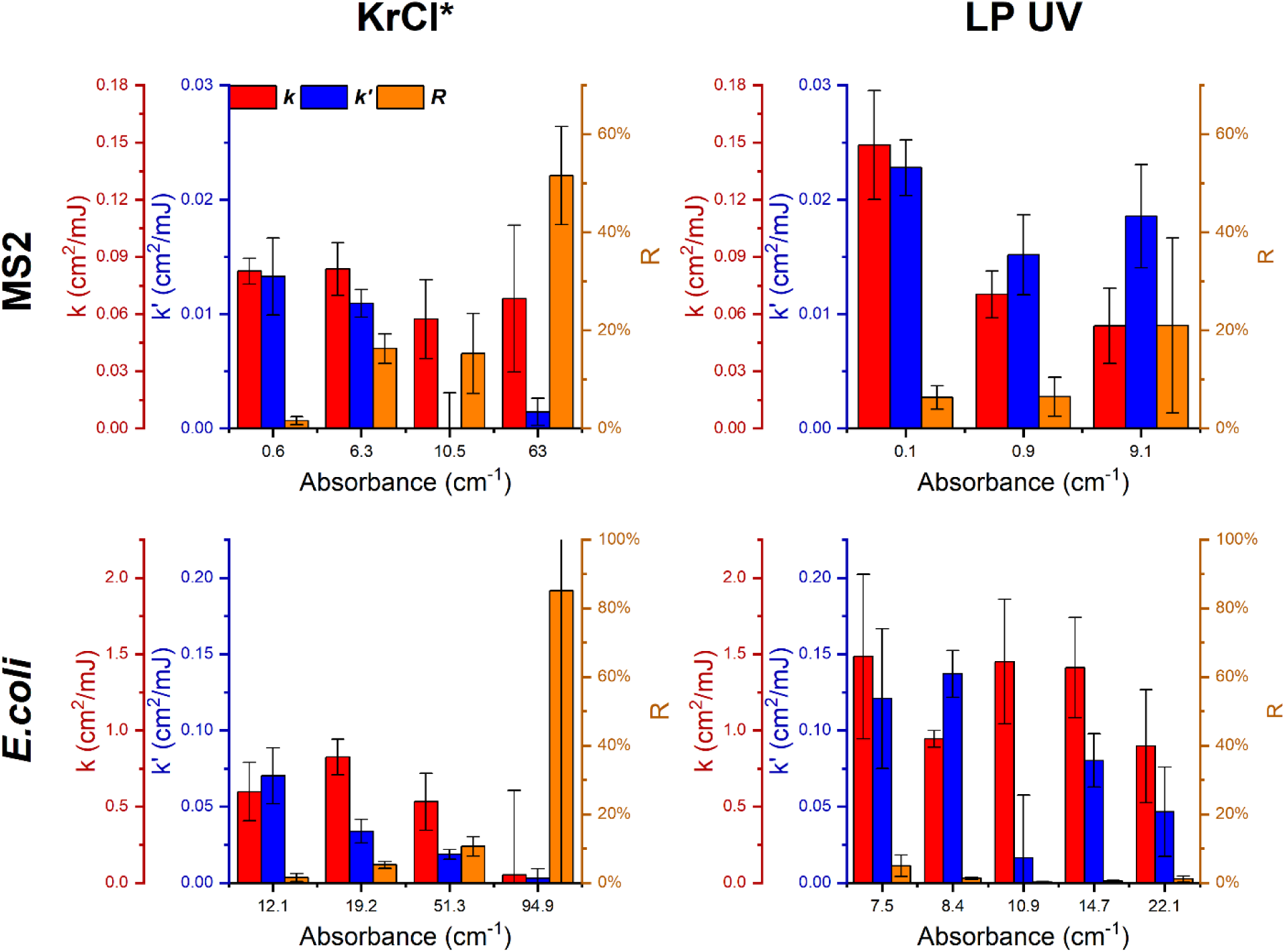
UV surface inactivation kinetics parameters for MS2 and *E. coli* using KrCl* excimer and LP UV lamp using the Spray-Swab method across deposition solution absorbance. The kinetics parameters were calculated using the ‘non-linear curve fitting’ function by fitting the *LRV* and *D* using Eq. 7. All UV inactivation results in this figure were obtained using the Spray-Swab method.

The reduction in UV inactivation performance with solution absorbance was greater for the KrCl* excimer than for the LP UV lamp (Fig. 2-A and B), indicating a greater physical shielding effect at 222 nm. This shielding effect from dried constituents (e.g., salts and proteins in the deposition solution) was likely achieved through two mechanisms: (1) small deposits that were developed on the surface of test microorganisms may absorb UV irradiation (Fig. 3: Mechanism A), and (2) large deposits (i.e., bigger than the microorganisms) may completely block or reflect/scatter the UV irradiation, depending on their relative location to the microorganisms (i.e., atop or adjacent to the microorganisms; Fig. 3: Mechanisms B and C). The greater reduction in inactivation performance using KrCl* excimer was likely attributed to the greater UV absorbance of the deposition solution at 222 nm compared to 254 nm (Fig. 2-C and D). Other mechanisms, such as UV blockage and reflection/scattering, are expected to be comparable at 222 nm and 254 nm, given that the same deposition solutions were used, resulting in similar deposit sizes and quantities on surfaces, and UV reflectance tends to be consistent across these wavelengths^41^. These findings suggest that UV absorption is likely the primary protective mechanism in UV surface disinfection. Future investigations on characterization of dried constituents on surfaces, such as chemical composition of the deposition solution, UV scattering and absorption profile, dried deposit morphology and spatial distribution of microorganisms within dried constituents, are needed to confirm the roles of different shielding mechanisms.

### Comparing UV sensitivities of microorganisms on surfaces and in aqueous solutions using KrCl* excimer and LP UV

Both KrCl* excimer and LP UV lamp can inactivate all test microorganisms on surfaces inoculated with solutions showing similar absorbance at 222 nm and 254 nm (Fig. 5 and Table 2). The absorbance values were within the range reported for human saliva (i.e., 50^th^ to 90^th^ percentile UV absorbance at 222 nm and 254 nm: 8-12.5 cm^-1^ and 6.2-12.2 cm^-1^, respectively^32^). Similar performance of KrCl* excimer and LP UV lamp were observed for bacteria (*E. coli* and *S. aureus*, paired t tests: P> 0.05; Table S3), whereas significantly higher inactivation of bacteriophage (MS2 and Phi6) was achieved using LP UV lamp compared to KrCl* excimer (Paired t tests: P< 0.05; Table S3). These results suggest that conventional LP UV lamp likely provides better surface disinfection performance for the same UV dose applied. However, KrCl* excimer was also very effective for disinfection and remains a compelling option, especially in occupied spaces, considering 222 nm UV irradiation is much safer than 254 nm (i.e., 25 and 45 times safer for human eye and skin exposures, according to American Conference of Governmental Industrial Hygienists (ACGIH) threshold limit values^42^).

**Figure 5.**
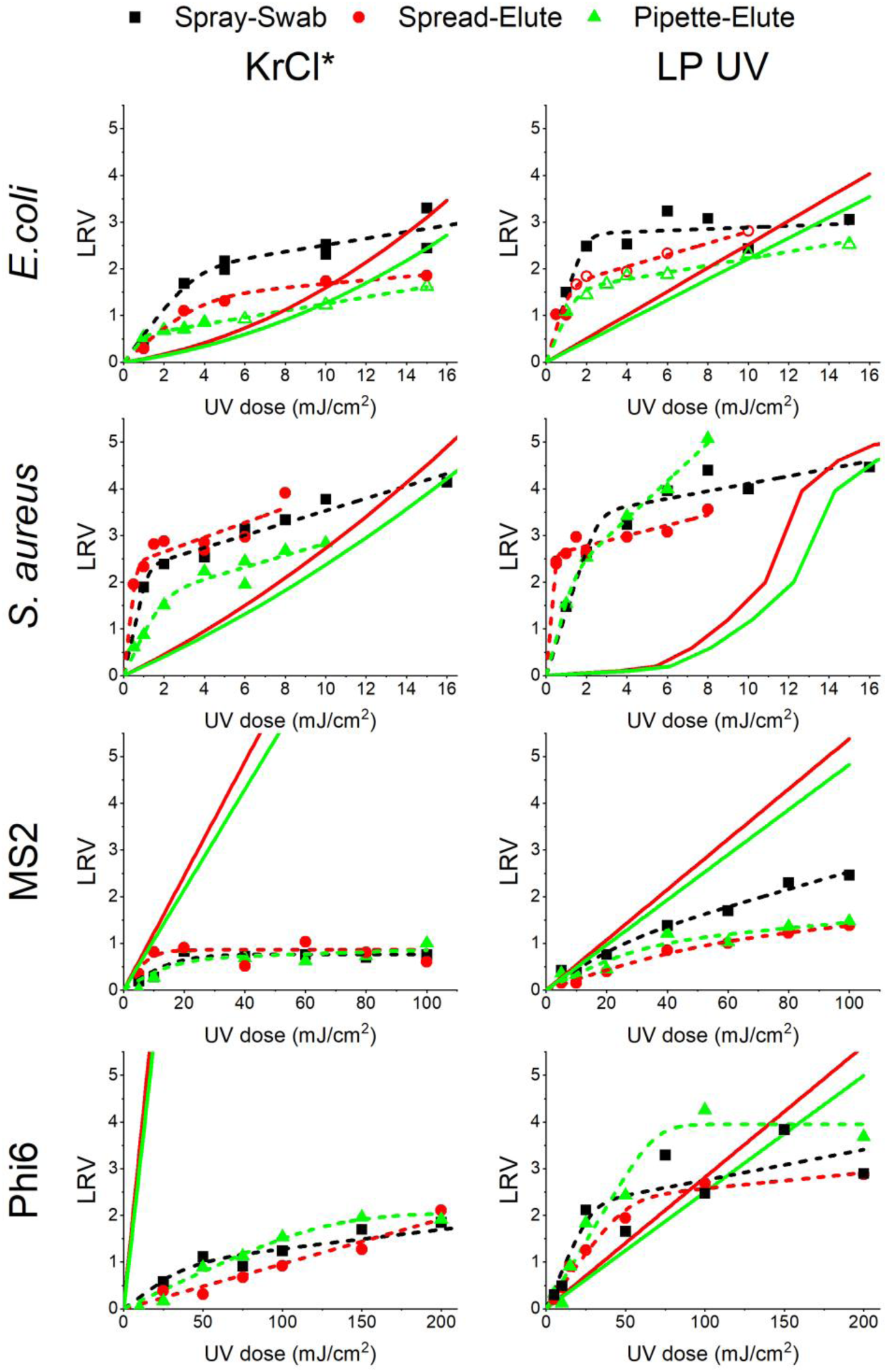
Comparison of UV surface inactivation tests using method proposed in this study and ASTM methods (Spray-Swab method: squares; Spread-Elute method: circles; Pipette-Elute method: triangles). Dashed lines represent the non-linear regression models computed based on Eq. 7. Solid lines represent the estimated UV dose response of the microorganisms in the deposition solutions used in the Spread-Elute method and the Pipette-Elute method before drying on surfaces, as described in Method Section 2.6.

Different trends in the UV dose response were observed for bacteria and viruses comparing surface and simulated aqueous conditions. In aqueous solutions, 222 nm UV inactivation is more effective for viruses but less effective for bacteria than 254 nm due to greater viral protein damage but less penetration into bacterial cellular components (cell wall, plasma membrane, and cytoplasm matrix)^18^. These trends, however, are opposite on surfaces as discussed above (Fig. 5). Comparing the UV dose response on surfaces and in aqueous solutions, higher UV inactivation of bacteria (*E. coli* and *S. aureus*) was observed on surfaces, especially at lower UV doses, whereas viruses exhibited greater UV resistance on surfaces (Fig. 5). These findings are consistent with those reported by Ratliff et al.^22^, which greater UV inactivation of MS2 was observed on wet surfaces than dried surfaces. The study by Duering et al., however, suggested greater UV inactivation of bacteria in aqueous solutions than on surfaces, likely due to a much higher UV dose applied in the inactivation test. Note that greater inactivation of bacteria in aqueous solutions was also observed in this study at higher UV doses (e.g., 16 mJ/cm^2^ in Fig. 5).

One possible explanation for the different bacterial UV dose response behavior on surfaces and in aqueous solution is cell desiccation. Bacteria cells may be desiccated on surfaces considering severe osmotic stresses from increasing salt and protein concentrations outside the cell membrane during evaporation^43^. Desiccation may induce structural changes in bacteria cells and macromolecules such as DNA and protein^44–46^, which may result in greater UV sensitivities compared to in an osmotically balanced state. Future investigations are needed to understand the detailed mechanisms on how cell desiccation affects bacterial UV sensitivities. Note that any desiccation-induced decay during the 1-hour drying period is not included in the inactivation results shown in Fig. 5. All control samples (no UV exposure) were collected after the 1-hour drying period and throughout the UV exposure in each experiment (see method section for details). No significant changes in bacterial/viral concentrations were observed in the triplicate control samples, indicating that the microbial viability did not change significantly across the UV exposure period (normally less than 1 hour). At higher UV doses, bacterial inactivation on surfaces was similar or even lower than in solutions (Fig. 5), likely due to the physical shielding effects provided by the dried deposits, especially in the deeper “UV resistant” regions (Fig. 3), which can protect a portion of the cells from UV exposure.

Unlike bacteria, viruses tend to maintain structural stability under severe osmotic conditions from tight genome packaging and lower internal water content^44,47^, meaning virus desiccation unlikely leads to a significant increase in the UV sensitivity in the way bacteria may be affected. Moreover, viruses are likely more susceptible to shielding by dried deposits because of their smaller size, which increases the extent of UV irradiation being absorbed or blocked by the deposits before reaching the viruses. This protective effect could be greater at lower wavelengths, as greater viral inactivation was observed at 254 nm than 222 nm, even with similar deposition solution absorbance at respective wavelengths (Fig. 5). This is further supported by the regression analysis, which the inactivation rate constants in the resistant region (*k’*) are consistently lower for KrCl* excimer than LP UV lamp for all test microorganisms (Table S1), indicating greater difference in the protective effect in the deeper UV resistant region (Fig. 3). One possible explanation is that UV light may be absorbed differently in salt/protein crystals compared to in aqueous solutions, and the UV absorbance measured in the deposition solutions may not accurately represent the extent of UV absorption by the salt/protein crystals across wavelengths.

### Comparing methods for UV surface disinfection tests

Minor differences in UV dose response curves were observed across surface disinfection methods, and the discrepancies varied with test microorganisms (Fig. 5). Comparing the three testing methods, the LRV achieved using the Spray-Swab method was marginally higher than using the Pipette-Elute method, especially for bacteria, with p values of less than 0.05 for 3 out of 4 testing conditions (paired t tests. *E. coli* and *S. aureus* using KrCl*, *E. coli* using LP; Fig. 5 and Table S3). The Spread-Elute method, however, exhibited no significant difference in UV inactivation to the other two methods in most testing conditions (P values > 0.05 for 6 out of 8 paired t tests). The differences in UV inactivation were likely due to the different deposition methods, which may result in different thickness and sizes of the droplets deposited on surfaces. The average deposit thickness was estimated as 0.055 mm and 0.164 mm for the Spread method and Pipette method, respectively (see detailed calculation in SI, Equation SI-5). While the average thickness for the Spray method could not be determined accurately, the observed surface coverage suggests it was similar to the Pipette method and thicker than the Spread method (see detailed calculation in SI). These results suggest that difference in deposit thickness across methods had limited impact on the UV inactivation efficacies. This is not consistent with previous studies that investigated UV inactivation by spreading or pipetting microorganisms on surfaces which found that spreading resulted in greater inactivation^22,29,30^. One possible explanation for this discrepancy is the higher UV absorbance of the deposition solutions used in the previous studies, which may amplify the effect of deposit thickness (i.e., more deposits accumulated per thickness due to higher concentrations of constituents) and result in more significant physical shielding effects. The droplet size, however, may better explain the differences in UV inactivation between the Spray and Pipette methods. Most droplets deposited using the Spray method were observed to be much smaller than the 1 µL droplets using the Pipette method (Fig. S2). Considering “coffee-ring” structure formation requires a minimum droplet size^48^, it is likely that there was less “coffee-ring” structures formed using the Spray method, resulting in less physical shielding effect and greater UV inactivation, as observed in Figure 5. Future investigations are needed to confirm the effect of “coffee-ring” structures on UV dose response on surfaces across droplet sizes.

Similar UV inactivation was observed for bacteriophage across methods (MS2 and Phi 6, Fig. 5), indicating that variations in thickness and droplet sizes from different deposition methods had limited impact on viral inactivation. Compared to bacterial cells, bacteriophage have much smaller sizes (23-28 nm for MS2^49^ and 85 nm for Phi 6^50^ vs. > 1 µm for *E. coli* and *S. aureus*^51,52^), resulting in greater physical shielding effects by the salt/protein deposits via UV light blockage. Thus, only the bacteriophages located at or near the surface of the dried deposits can be effectively exposed to UV. This is further supported by the lower maximum LRV observed for bacteriophage compared to bacteria (Fig. 5). These results suggest that the impact of surface deposition on UV disinfection efficacy varies with microorganisms and should be taken into consideration in designing experiments to evaluate UV disinfection performance on surfaces.

Table 3 summarizes the advantages and disadvantages of all deposition methods evaluated in this study. Compared to the standard ASTM methods (i.e., the Spread and Pipette method^35,36^), the main advantage of the Spray method is that it produces microorganism-containing droplets with variable sizes and thicknesses, which may better approximate real-world pathogen deposition scenarios such as sneezing or coughing on surfaces^53,54^. A major limitation of the Spray method, however, is the potential variability between tests due to spray bottle selection and operation, which may lead to inconsistency in UV inactivation results due to different droplet sizes and thickness on surfaces. Similar issues may also occur using the Spread methods, considering evenly spreading deposition solutions on surfaces may be difficult, especially for hydrophobic materials due to lower surface energy^55^. The Pipette method is relatively easy to perform with more consistent droplet size and shape. However, the droplets produced tend to be much bigger than those in the real-world scenarios (droplet diameter: 1.2 mm for 1 µL droplet vs. less than 0.38 mm in sneeze and coughing^53,54^), providing more physical shielding effects and underestimating the UV disinfection performance, especially for emerging far UV devices like KrCl* excimer lamps. While smaller droplets were observed using the Spray method (Fig. S2), it is uncertain that their size distribution accurately represents real-world pathogen deposition due to the lack of quantitative characterization. Overall, while each deposition method has strengths and limitation, all methods can benefit from improved consistency and similarity to real-world deposition conditions in the droplet production.

**Table 3.** The advantages and disadvantages of all deposition and extraction methods used in this study.

|  | Advantages | Disadvantages |
| --- | --- | --- |
| **Deposition methods** |  |  |
| **Spray** | Variable droplet sizes and thickness | Potential inconsistency between tests |
| **Spread** | Similar thickness to real droplet | Difficult to spread on hydrophobic surfaces |
| **Pipette** | Greater consistency | Underestimating UV disinfection performance due to greater droplet thickness |
| **Extraction methods** |  |  |
| **Swab** | Greater extraction efficiency and consistency<br>Greater LRV detection limit | Potential poor performance for porous materials |
| **Elute** | Easy to perform | Lower extraction efficiency<br>Lower LRV detection limit |

The extraction methods unlikely affected the UV inactivation results, considering both control samples (i.e., no UV exposures) and samples post UV exposures were extracted using the same method and any differences in extraction methods were cancelled out in the LRV calculations. Among tested microorganisms, *E. coli* exhibited significantly lower extraction efficiency compared to others (Table 2). This is because *E. coli* cells likely decay much faster during the 1-hour drying period^56,57^, which leads to a lower calculated extraction efficiency considering it was calculated by comparing the amount of microorganisms deposited on surfaces before drying vs. the amount recovered after drying and extraction. The LRV detection limits were vastly different across extraction methods due to the differences in the volume of extraction solution used and the extraction efficiency (Table 2). Compared to the Elute method, the Swab method provided higher extraction efficiency for bacteria and comparable extraction efficiency for viruses (Table 2). This is likely due to more effective physical removal of the microorganisms from surfaces via swabbing than eluting using the extraction solution, despite that some recovered microorganisms may be retained on the cotton swab rather than transferred into the extraction solution. Combining higher extraction efficiencies and lower volume of extraction solution (Swab vs. Elute: 1 mL vs. 20 mL), the Swab method provided much higher LRV detection limits, typically greater by a LRV of 2 than the Elute method (Table 2). In addition, the extraction efficiencies using the Swab method were observed to be more consistent than the Elute method, as indicated by lower relative standard errors (RSE) for the extraction efficiencies (Table 2). Considering both the LRV detection limit and the extraction performance consistency (Table 3), the Swab method is recommended for microorganism extraction in UV surface disinfection tests. A potential limitation of the Swab method, however, is its reduced performance on porous or highly textured materials, which microorganisms may penetrate below the surface and be difficult to recover (Table 3). Additionally, both Swab and Elute methods exhibited noticeable variations in extraction performance (RSE in Table 2), likely attributed to the inconsistencies associated with manual extraction operation and other operator-induced errors. Future investigations are needed to evaluate the performance of the Swab method on other materials and explore automated extraction methods to improve the extraction performance consistency.

Overall, this study demonstrates that germicidal UV devices can effectively inactivate bacteria and viruses on surfaces. The observed UV dose response on surfaces followed a two-region non-linear kinetics due to the physical shielding effects from the dried deposition constituents. Absorption was found to be the primary mechanism of the shielding effect, with other mechanisms such as reflection and blockage also playing an important role. On surfaces inoculated with solutions having similar UV absorbance as human saliva, LP UV lamp exhibited similar bacterial inactivation and greater viral inactivation compared to KrCl* excimer lamp at the same UV dose. However, KrCl* could be more favorable, especially in occupied spaces, because of its relative safety, which allows a much higher UV dose to be applied. Comparison between surface and aqueous inactivation revealed that cell desiccation on surfaces may enhance bacterial UV sensitivity, while viruses remained more resistant due to structural stability and greater susceptibility to shielding due to smaller size. One limitation of this study is that all experiments were conducted on polycarbonate surfaces under uncontrolled RH. The UV disinfection performance across wavelengths, relative roles of shielding mechanisms, and microbial UV sensitivities may vary across other surface materials and RH. UV inactivation on surfaces also varied with deposition methods (i.e., Spray, Spread, and Pipette), which the Spray method achieved higher inactivation, especially for bacteria, due to less physical shielding from smaller droplet sizes. While both extraction methods (i.e., Swab and Elute) did not affect the inactivation, the Swab method is recommended due to higher LRV detection limits and more consistent extraction efficiency. In sum, these findings emphasize the importance of deposition conditions (absorbance and method), UV wavelengths (222 nm vs. 254 nm), and microbial characteristics (bacteria vs. viruses) on UV surface disinfection, and provide guidance on future design and testing procedures of germicidal UV devices for effective surface disinfection, especially in high-risk occupied spaces.

## ACKNOWLEDGMENTS

Financial support for this work was provided by the National Science Foundation, Grant CBET 2029695. We thank Dr. Michael Fisher and Emanuele Sozzi for providing bacteriophage Phi6 and *Pseudomonas syringae*.

