## Supplemental Information for "UV inactivation of bacteria and viruses on surfaces: mechanistic insights and testing method comparisons"

### Supporting Information

Ben Ma[a,b,*], Saba Seyedi[a,c], and Karl G. Linden[a,*]

a. Department of Civil, Environmental, and Architectural Engineering, University of Colorado Boulder, 4001 Discovery Dr., Boulder, CO, 80303, United States
b. Department of Civil and Environmental Engineering, University of Nevada, Reno, 1664 N Virginia St., Reno, NV, 89557, United States
c. Jacobs Engineering, 1851 Alexander Bell Drive, Reston, VA, 20191, United States

*Corresponding authors:

### Methods for bacteria and virus stock preparation and quantification

#### MS2 coliphage

The method for MS2 propagation was adopted from standard method 1601 of US EPA (7). The *E. coli* $F_{amp}$ host was grown in Tryptic Soy Broth (TSB) with streptomycin and ampicillin at 37 °C and 200 rpm for 2-4 hours to an optical density ($OD_{600}$) of 0.1-1.0 at 600 nm. Then, 5 mL cell culture and 20 uL MS2 stock was added into a propagation mixture containing 50 mL sterile DI water, 50 mL TSB, 1.25 mL 4M $MgCl_2 \cdot 6H_2O$ solution, and 0.5 mL streptomycin ampicillin solution (3 g/L for each antibiotics) and then incubated under the same conditions for another 18-24 hours. The propagation mixture was then centrifuged at 4000 g for 30 min, and the supernatant was transferred and filtered through 0.22 µm polyethersulfone membranes to remove cells and debris. The infectivity of MS2 was measured by plaque assay on *E. coli* $F_{amp}$ cells following a two layer agar procedure (standard EPA method 1601 (7)). Briefly, bottom layer agar plates were prepared by dispensing 15mL 1% Tryptic Soy Agar (TSA; containing 30 g/L TSB and 10 g/L Bacto™ agar). Top layer agar mixture was prepared on the day of plaque assay, containing 5 mL 0.75% TSA (30 g/L TSB and 7.5 g/L Bacto™ agar) and 0.5 mL *E. coli* $F_{amp}$ cell culture with $OD_{600}$ of 0.1-1.0. Serial 10-fold dilutions of MS2 samples were prepared in 0.01M phosphate buffered saline (PBS; pH 7.4), and 0.1 mL of dilution was added into the top layer agar mixture. The top layer agar mixture was then dispensed onto the bottom layer agar plates. The agar plates were incubated at 37 °C for 18 to 24 hours and then the plaque-forming units (PFU) were counted. At least three consecutive dilutions were plated and the plates with PFU of 0-200 were recorded. The viable MS2 concentration was calculated as:

$$C = \frac{N_1 + N_2 + N_3}{10^{-D_1} + 10^{-D_2} + 10^{-D_1}} \cdot \frac{1}{0.1\,mL}$$

where $C$ is the MS2 concentration in PFU/mL, $N_1$ $N_2$ $N_3$ are the PFU numbers per plate (0-200) at dilutions $D_1$, $D_2$, and $D_3$, respectively.

**Phi 6.**

Bacteriophage Phi 6 and its bacterial host *Pseudomonas syringae* were kindly provided by Dr. Michael Fisher's group at University of North Carolina at Chapel Hill. The method for Phi6 propagation was adopted from standard method 1602 of US EPA[1]. *P. syringae* was grown in Tryptic Soy Broth (TSB) at room temperature (25 ºC) and 140 rpm for 18-24 hours to an optical density ($OD_{600}$) of 0.1-1.0 at 600 nm. Then, 5 mL *P. syringae* cell culture and 20 µL Phi 6 stock was added into a propagation mixture containing 50 mL sterile DI water, 50 mL TSB, and 1.25 mL 4M $MgCl_2 \cdot 6H_2O$ solution, and then incubated under the same conditions for another 18-24 hours. The propagation mixture was then centrifuged at 4000 g for 30 min, and the supernatant was transferred and filtered through 0.22 µm polyethersulfone membranes to remove cells and debris. The infectivity of Phi6 samples was measured by plaque assay on *P. syringae* cells following a two layer agar procedure (standard EPA method 1601[2]). Briefly, bottom layer agar plates were prepared by dispensing 15mL 1% Tryptic Soy Agar (TSA; containing 30 g/L TSB and 10 g/L Bacto™ agar). Top layer agar mixture was prepared on the day of plaque assay, containing 5 mL 0.75% TSA (30 g/L TSB and 7.5 g/L Bacto™ agar) and 0.5 mL *P. syringae* cell culture with $OD_{600}$ of 0.1-1.0. Serial 10-fold dilutions of Phi 6 samples were prepared in 0.01M phosphate buffered saline (PBS; pH 7.4), and 0.1 mL of dilution was added into the top layer agar mixture. The top layer agar mixture was then dispensed onto the bottom layer agar plates. The agar plates were incubated at room temperature for 18 to 24 hours and then the PFU were counted for at least three dilutions. The viable Phi6 concentration was calculated using the same equation for MS2 as described above.

***Escherichia coli K-12.***

Stationary phase cultures of *Escherichia coli K-12* (ATCC 29425) was grown from frozen stored stocks in Luria-Bertani (LB) broth and incubated for 16-18 hours at 37°C and shaking at 180 rpm (1). To prepare the test water, the overnight culture was pelletized by centrifugation (3000 × *g*, 5 minutes). The supernatant was collected in a different sterile conical tube and the remaining pellet was resuspended in PBS (~$10^9$ CFU/mL). This was repeated three times to "wash" the original growth media from the cells. The viable *E. coli* cell concentration was determined using a standard spot plate counting method. Briefly, serial 10-fold dilutions of samples were prepared in PBS, and triplicate 10 µL of six consecutive dilutions were plated on LB agar plates as spots. The agar plates were incubated at 37 °C for 16-20 hours and then the colony forming units (CFU) were counted. Only the spots with CFU of 0-50 were recorded and the concentration was calculated based on the log-scale average of triplicate plating as:

$$\log C = averager \left[\log\left(\frac{N_1}{10^{-D_1}}\right), \log\left(\frac{N_2}{10^{-D_2}}\right), \log\left(\frac{N_3}{10^{-D_3}}\right)\right] + \log\left(\frac{1}{0.01\ mL}\right)$$

where $C$ is the *E.coli* concentration in CFU/mL, $N_1$ $N_2$ $N_3$ are the CFU numbers per spot at dilutions $D_1$, $D_2$, and $D_3$, respectively, counted from the triplicate plating.

***Staphylococcus aureus***

*Staphylococcus aureus* (ATCC 6538) were grown from frozen stored stocks in sterile tryptic soy broth (TSB) and incubated for 16-18 hours at 37°C and shaking at 200 rpm (4). To prepare the test water, the overnight culture was pelletized by centrifugation (3000 × *g*, 5 minutes). The supernatant was discarded and the remaining pellet was resuspended in PBS (~$10^9$ CFU/mL). This was repeated three times to "wash" the original growth media from the cells. For *S. aureus* cell cultures, after the pellet is resuspended the third time, the sample was sonicated for 60 seconds at

82 Watts to disaggregate the cells. The viable *S. aureus* concentrations were determined on tryptic soy agar plate using the standard spot plate counting method as described above.

**Transformation of Eq.7 from Geeraerd and Van Impe inactivation model:**

Original Geeraerd and Van Impe inactivation model:

$$N = (N_0 - N_r) \times 10^{-kD} + N_r \qquad SI-1$$

Two-region inactivation model adopted:

$$N = (N_0 - N_r) \times 10^{-kD} + N_r \times 10^{-rkD} \qquad SI-2$$

Equation divided by $N_0$ on both side:

$$\frac{N}{N_0} = \left(1 - \frac{N_r}{N_0}\right) \times 10^{-kD} + \frac{N_r}{N_0} \times 10^{-rkD} \qquad SI-3$$

Thus,

$$LRV = log_{10}\left(\frac{N_0}{N}\right) = -log_{10}\left[\left(1 - \frac{N_r}{N_0}\right) \times 10^{-kD} + \frac{N_r}{N_0}\right]$$

$$= -\log[(1-R) \times 10^{-kD} + R \times 10^{-rkD}] \qquad SI-4$$

where $N_r$ is the number of microorganisms that are resistant to UVC irradiation, *D* is UV fluence in mJ/cm$^2$ between 200 nm to 400 nm, *k* is the pseudo-first-order inactivation rate constant (in cm$^2$/mJ) for the microorganisms in the inactivation region, *rk* is the pseudo-first-order inactivation rate constant (in cm$^2$/mJ) for the microorganisms in the resistant region, which *r* (≤ 1) is the resistant factor for the inactivation rate constant, and *R* is the fraction of the microorganisms in the resistant region.

**Estimation of the average thickness of deposited solution on surfaces**

The average thickness of the solution deposited on surfaces was estimated using the following equation:

$$Average\ thickness = \frac{Deposition\ volume}{Surface\ area} \qquad SI-5$$

For the Spread method, the deposition volume is 20 μL ($mm^3$) and surface area is 19 mm×19 mm =316 $mm^2$. The average thickness was calculated as **0.055** mm.

For the Pipette method, the deposition volume is 1 μL ($mm^3$) for each droplet. The diameters of 10 droplets were measured, with the average diameter (± standard deviation) of 2.79 ± 0.46 mm. Thus, the surface area of each droplet was estimated as $\frac{\pi}{4} \times (2.79\ mm)^2 = 6.11\ mm^2$, and the average thickness was calculated as **0.164** mm.

Considering the surface area of the deposition solution using the Spray method cannot be easily measured, the average thickness was estimated across the coverage of the inoculation solutions on the surface coupons using the following equation as shown in Fig. S1:

$$Average\ thickness = \frac{Deposition\ volume}{Coupon\ surface\ area \times Coverage\ in\ \%} \qquad SI-6$$

where the coupon surface area is 1963.5 $mm^2$ (coupon diameter = 50 mm).

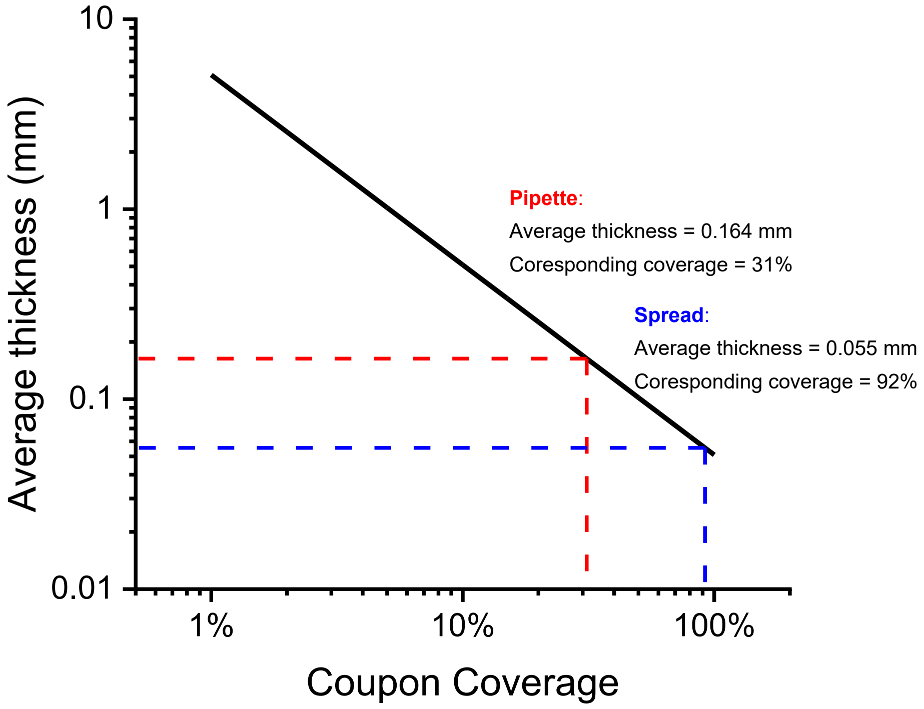


**Figure S1.** Average thickness of deposition solution for Spray-Swab method. The deposition volume of 100 µL is used for the calculation.

Using the relationship between the average thickness and the coupon coverage, the corresponding coverage to achieve the same thickness as Spread and Pipette methods were determined as 92% and 31%, respectively. Based on the observations of surface coupons right after inoculation (Fig. S2), The Spray method likely provided an average thickness similar to the Pipette method but greater than the Spread method based on the observed coverage (20-50 %)

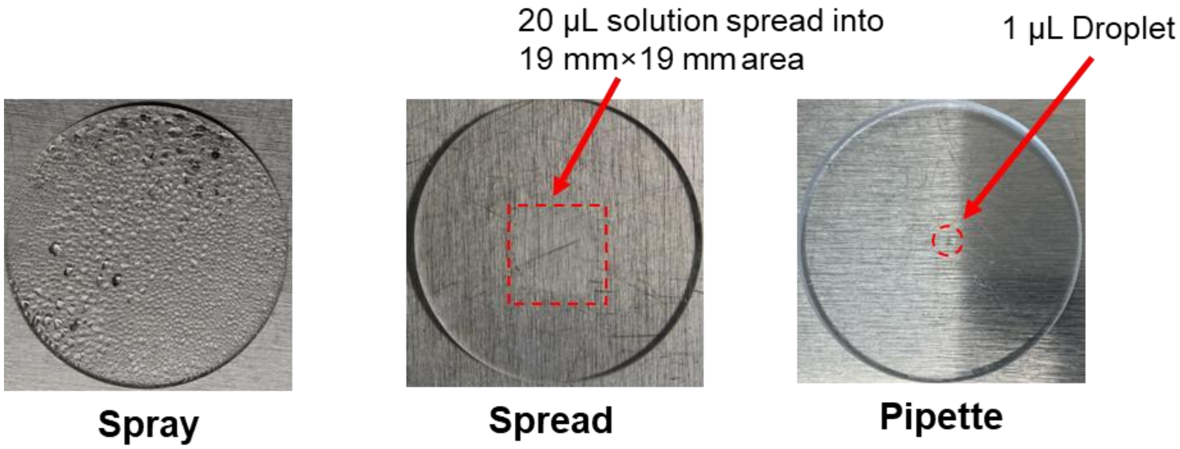


**Figure S2.** Pictures of coupons right after solution deposition.

**Table S1.** Results of non-linear regression model for UV dose response on surfaces using KrCl* excimer and LP UV lamp.

| **Inoculation solution** | **Testing Method** | **KrCl* excimer (222 nm)** | | | | **LP UV (254 nm)** | | | |
|---|---|---|---|---|---|---|---|---|---|
| | | $k$ (cm$^2$/mJ) | $k'$ (cm$^2$/mJ) | R (in %) | $R^{2\Delta}$ | $k$ (cm$^2$/mJ) | $k'$ (cm$^2$/mJ) | R (in %) | $R^2$ |
| **MS2** | | | | | | | | | |
| 100-fold diluted | 1 | 0.082 ± 0.007 | 0.013 ± 0.003 | 1.6 ± 0.8 | 0.97 | 0.149 ± 0.029 | 0.023 ± 0.002 | 6.3 ± 2.4 | 0.99 |
| 10-fold diluted | 1 | 0.084 ± 0.014 | 0.011 ± 0.001 | 16.3 ± 3.0 | 0.98 | 0.070 ± 0.012 | 0.015 ± 0.004 | 6.5 ± 4.0 | 0.98 |
| Non-diluted | 1 | 0.068 ± 0.038 | 0.001 ± 0.001 | 51.6 ± 10.0 | 0.83 | 0.054 ± 0.020 | 0.019 ± 0.005 | 21.0 ± 17.8 | 0.98 |
| For KrCl*: 6-fold diluted For LP UV: Non-diluted | 1 | 0.057 ± 0.021 | 0.000 ± 0.003 | 15.3 ± 8.1 | 0.82 | 0.054 ± 0.020 | 0.019 ± 0.005 | 21.0 ± 17.8 | 0.98 |
| | 2 | 0.118 ± 0.071 | 0.000 ± 0.004 | 12.6 ± 8.4 | 0.18 | 0.026 ± 0.007 | 0.006 ± 0.005 | 15.1 ± 18.0 | 0.99 |
| | 3 | 0.057 ± 0.044 | 0.002 ± 0.005 | 22.6 ± 19.4 | 0.74 | 0.040 ± 0.020 | 0.006 ± 0.007 | 13.5 ± 17.7 | 0.91 |
| ***E. coli* K-12** | | | | | | | | | |
| Resuspended in PBS | 1 | 0.598 ± 0.192 | 0.070 ± 0.018 | 1.6 ± 1.1 | 0.90 | 1.484 ± 0.540 | 0.121 ± 0.046 | 5.0 ± 3.1 | 0.84 |
| Re-suspended in 10-fold diluted supernatant | 1 | 0.826 ± 0.117 | 0.034 ± 0.008 | 5.2 ± 1.0 | 0.96 | 0.944 ± 0.055 | 0.137 ± 0.015 | 1.3 ± 0.4 | 1.00 |
| Re-suspended in 2-fold diluted supernatant | 1 | 0.532 ± 0.187 | 0.019 ± 0.003 | 10.6 ± 2.8 | 0.95 | 1.411 ± 0.329 | 0.080 ± 0.017 | 0.5 ± 0.2 | 0.96 |
| Re-suspended in undiluted supernatant | 1 | 0.052 ± 0.554 | 0.003 ± 0.006 | 85.2 ± 93.8 | 0.42 | 0.896 ± 0.371 | 0.047 ± 0.029 | 1.1 ± 0.9 | 0.72 |
| For KrCl*: Resuspended in PBS For LP UV: Re-suspended in 5-fold diluted supernatant | 1 | 0.598 ± 0.192 | 0.070 ± 0.018 | 1.6 ± 1.1 | 0.90 | 1.450 ± 0.408 | 0.016 ± 0.041 | 0.2 ± 0.2 | 0.76 |
| | 2 | 0.404 ± 0.073 | 0.038 ± 0.022 | 5.0 ± 3.1 | 0.99 | 1.516 ± 0.295 | 0.125 ± 0.026 | 2.8 ± 1.1 | 0.98 |
| | 3 | 1.268 ± 0.316 | 0.073 ± 0.003 | 30.2 ± 1.8 | 1.00 | 1.138 ± 0.159 | 0.074 ± 0.009 | 3.3 ± 0.6 | 0.98 |
| ***Bacteriophage Phi6*** | | | | | | | | | |
| For KrCl*: 9-fold diluted For LP UV: 1.75-fold diluted | 1 | 0.032 ± 0.016 | 0.004 ± 0.002 | 13.3 ± 8.6 | 0.86 | 0.082 ± 0.037 | 0.007 ± 0.005 | 0.8 ± 1.3 | 0.81 |
| | 2 | 0.010 ± 0.001 | 0.000 ± 0.000 | 0.0 ± 35.8 | 0.95 | 0.048 ± 0.004 | 0.003 ± 0.002 | 0.6 ± 0.4 | 0.99 |
| | 3 | 0.016 ± 0.003 | 0.000 ± 0.007 | 0.9 ± 2.8 | 0.97 | 0.057 ± 0.008 | 0.000 ± 0.006 | 0.0 ± 0.0 | 0.95 |
| ***S. aureus*** | | | | | | | | | |
| For KrCl*: resuspended in PBS then diluted 3.5-fold For LP UV: resuspended in PBS then diluted 2.5-fold | 1 | 2.064 ± 0.531 | 0.132 ± 0.018 | 0.6 ± 0.2 | 0.96 | 1.373 ± 0.257 | 0.080 ± 0.035 | 0.0 ± 0.0 | 0.94 |
| | 2 | 3.797 ± 2.050 | 0.157 ± 0.049 | 0.5 ± 0.2 | 0.82 | 5.716 ± 1.467 | 0.110 ± 0.025 | 0.3 ± 0.1 | 0.89 |
| | 3 | 0.955 ± 0.212 | 0.127 ± 0.045 | 2.8 ± 2.0 | 0.96 | 1.686 ± 0.328 | 0.403 ± 0.040 | 1.8 ± 1.0 | 1.00 |

$^{\Delta}$ Regression correlation coefficient.

**Table S2.** Kendall's correlation between the deposition solution absorbance and UV surface inactivation kinetics parameters (*k*, *k'*, and *R*) for MS2 and *E. coli*.

| | **MS2-KrCl** | | **MS2-LP** | | ***E. coli*-KrCl** | | ***E. coli*-LP** | |
|---|---|---|---|---|---|---|---|---|
| | **τ** | **P-value** | **τ** | **P-value** | **τ** | **P-value** | **τ** | **P-value** |
| *k* | -0.33 | 0.50 | -1.00 | 0.00* | -0.67 | 0.17 | -0.60 | 0.14 |
| *k'* | -0.67 | 0.17 | -0.33 | 0.60 | -1.00 | 0.00* | -0.40 | 0.33 |
| *R* | 0.67 | 0.17 | 1.00 | 0.00* | 1.00 | 0.00* | -0.40 | 0.33 |

**Table S3.** P values of paired t tests comparing the results in Figure 5.

| UV device | KrCl* | | | LP | | | KrCl* vs. LP | | |
|---|---|---|---|---|---|---|---|---|---|
| **Method$^{+}$** | **1 vs. 2** | **1 vs. 3** | **2 vs. 3** | **1 vs. 2** | **1 vs. 3** | **2 vs. 3** | **1** | **2** | **3** |
| *E. coli* | 0.012$^{\Delta}$ (M1>M2) | 0.082$^{\Delta}$ (M1>M3) | 0.275 | 0.106 | 0.013$^{\Delta}$ (M1>M3) | 0.072 | 0.324 | 0.123 | 0.000$^{\Delta}$ |
| *S. aureus* | 0.080 | 0.001$^{\Delta}$ (M1>M3) | 0.001$^{\Delta}$ (M2>M3) | 0.662 | 0.292 | 0.489 | 0.062 | 0.460 | 0.010$^{\Delta}$ |
| MS2 | 0.272 | 0.883 | 0.370 | 0.004$^{\Delta}$ (M1>M2) | 0.024$^{\Delta}$ (M1>M3) | 0.015$^{\Delta}$ (M2<M3) | 0.033$^{\Delta}$ (KrCl*<LP) | 0.924 | 0.082 |
| Phi6 | 0.093 | 0.831 | 0.121 | 0.125 | 0.912 | 0.098 | 0.003$^{\Delta}$ (KrCl*<LP) | 0.008$^{\Delta}$ (KrCl*<LP) | 0.013$^{\Delta}$ (KrCl*<LP) |

+Method 1: Spray-Swab; Method 2: Spread-Elute; Method 3: Pipette-Elute.

$\Delta$ P value $< 0.05$
